# A survey of proteomic variation across two ethnic groups in Nigeria and its relationship to obesity risk

**DOI:** 10.1101/2022.12.09.519773

**Authors:** Arjun Biddanda, Karen Perez de Arce, Golibe Eze-Echesi, Chiamaka Nwuba, Yusuf Ibrahim, Olubukunola Oyedele, Esha Joshi, Boladale Alalade, Olanrewaju Ajayi, Chidimma Nwatu, Aminu Yakubu, Abasi Ene-Obong, Jumi Popoola, Colm O’Dushlaine, Peter Fekkes

**Affiliations:** 54gene, Inc. 1100 15th St NW, Washington, DC 20005, United States; Federal Medical Centre Abeokuta - Olabisi Onabajo way Idi-Aba, PMB 3031, Abeokuta, Ogun State, Nigeria; Federal Medical Centre Ebute-Metta - Railway Compound, PMB 1097 Ebute-Metta, Lagos, Nigeria; University of Nigeria Teaching Hospital - Ituku-Ozala, PMB 00129, Enugu, Enugu State, Nigeria

## Abstract

Proteomic variation between individuals has immense potential for identifying novel drug targets and disease mechanisms. However, with high-throughput proteomic technologies still in their infancy, they have largely been applied in large majority European ancestry cohorts (e.g. the UK Biobank). An open question is the degree to which proteomic signatures seen in European and other groups mirror those seen in diverse populations, such as cohorts from Africa. Coupled with genetic information, we can also gain a better understanding of the role of genetic variants in the regulation of the proteome and subsequent disease mechanisms.

To address the gap in our understanding of proteomic variation in individuals of African ancestry, we collected proteomic data from 176 individuals across two ethnic groups (Igbo and Yoruba) in Nigeria. These individuals were also stratified into high BMI (BMI > 30 kg/m^2^) and normal BMI (20 kg/m^2^ < BMI < 30 kg/m^2^) categories. We characterized differences in plasma protein abundance using the Olink Explore 1536 panel between high and normal BMI individuals, finding strong associations consistent with previously known signals in individuals of European descent. We additionally found 73 sentinel cis-pQTL in this dataset, with 21 lead cis-pQTL not observed in catalogs of variation from European-ancestry individuals. In summary, our study highlights the value of leveraging proteomic data in cohorts of diverse ancestry for investigating trait-specific mechanisms and discovering novel genetic regulators of the plasma proteome.

## INTRODUCTION

A key step towards understanding how genetic variation drives phenotypic change is a better understanding of how changes in DNA impact intermediate molecular traits. Molecular traits have substantially improved our understanding of the cascade of effects that genetic variants can have, and their contribution to larger organismal phenotypes [1,2]. One particularly attractive molecular measurement is to directly measure protein abundance, since proteins are a natural endpoint of how genetic information is translated from genes [3]. Recent advancements in targeted proteomics have allowed for richer characterizations of the proteome at larger scales. Insights into large-scale proteomic variation has led to improved understanding of causal mechanisms underlying disease, drug repurposing, and the impact of genetic variants on protein abundance [4–6].

Protein dysregulation can also function as a predictive biomarker for more complex traits, allowing actionable patient stratification. For example, levels of C-Reactive Protein (CRP) and fibrin-D have long been used as molecular biomarkers of rheumatic disease and cardiomyopathy, respectively [7–10]. Recent epidemiological evidence suggests that non-communicable diseases (NCDs) are presenting an increasing healthcare burden on the African continent, in contrast to the much larger investment in tackling communicable diseases over the past fifty years [11,12]. Therefore, larger proteomic surveys offer a potentially unique set of biomarkers to improve precision medicine and patient stratification efforts for NCDs in Africa.

Obesity is an increasingly prevalent health condition on the African continent, poised to be a considerable burden for healthcare systems over the coming decades. The genomics of obesity has been well-studied over the past decade, with large genomic databases now able to explain a sizable proportion of the variation in BMI (a proxy for obesity), and have led to a number of key drug targets [13,14]. In European ancestry individuals, proteomics has been instrumental in understanding causal links between genetic and proteomic factors in obesity pathogenesis [15]. For example, the role of *LEP/LEPR* in obesity has been well-established using bi-directional Mendelian randomization using cis-pQTL from European-ancestry individuals [15,16]. However, the integration of proteomic information for the study of obesity in individuals of African ancestry has been underrepresented in this body of literature.

To address these gaps in knowledge for individuals of African ancestry, we generated genetic and proteomic data from samples of individuals from two Nigerian ethnolinguistic groups (the Igbo and Yoruba). We selected individuals from these groups who were obese (defined as BMI > 30 kg/m^2^) as well as healthy controls (defined as BMI < 30 kg/m^2^) to identify differences in protein abundance underpinning obesity in African-ancestry populations, a primary aim of this study. A second aim of this study was to estimate the effect of *cis* genetic variants on protein abundance, with the hypothesis that ancestry-specific variants may be associated with protein levels. Taken together, these aims seek to both develop a proteomic resource for the study of obesity and understanding the impact of genetic variation on protein abundance in African-ancestry individuals.

## RESULTS

### Phenotype Overview

We collected phenotypic information from 171 individuals passing quality control (see Methods), largely centered around anthropometric measurements and baseline health (Table 1). As expected, many of the phenotypes that correlate with BMI are significantly different between the case and control groups (Table 1). These phenotypes are either components of BMI (e.g. weight) or are physiological or clinically correlated traits, such as systolic and diastolic blood pressure (Figure 1A). The correlation between these traits and our outcome variable of interest (obesity status) are important when performing subsequent regression analyses, to avoid introducing multi-collinearity and a reduction in statistical power. Outside of anthropometric traits and blood pressure, we note that most of the other phenotypes have correlation with BMI that are < 0.5 and are likely safer to include as covariates (Table1, Figure 1A).

**Table 1.** Epidemiological Description of Obesity Cases and Controls. Obesity cases and controls split across multiple phenotypes and sample-level metadata. P-values are adjusted for multiple testing using Bonferroni correction.

|  |  | Grouped by Disease_Code |  |  |  |  |
| --- | --- | --- | --- | --- | --- | --- |
|  |  | Missing | Overall | Case | Control | P-Value (adjusted) |
| **N** |  |  | 171 | 87 | 84 |  |
| **Site, n (%)** | **ABK** | 0 | 97 (56.7) | 45 (51.7) | 52 (61.9) | 1.000 |
|  | **EBM** | 0 | 46 (26.9) | 27 (31.0) | 19 (22.6) |  |
|  | **UNTH** | 0 | 28 (16.4) | 15 (17.2) | 13 (15.5) |  |
| **Age, mean (SD)** |  | 0 | 45.4 (12.6) | 45.9 (12.1) | 44.8 (13.1) | 1.000 |
| **Sex, N (%)** | **Female** | 0 | 120 (70.2) | 60 (69.0) | 60 (71.4) | 1.000 |
|  | **Male** | 0 | 51 (29.8) | 27 (31.0) | 24 (28.6) |  |
| **Ancestry (self-id), N (%)** | **Igbo** | 0 | 65 (38.0) | 34 (39.1) | 31 (36.9) | 1.000 |
|  | **Yoruba** | 0 | 106 (62.0) | 53 (60.9) | 53 (63.1) |  |
| **Weight (kg), mean (SD)** |  | 0 | 82.3 (25.4) | 101.8 (21.1) | 62.1 (6.9) | <0.001 |
| **Height (m), mean (SD)** |  | 0 | 1.7 (0.1) | 1.7 (0.1) | 1.7 (0.1) | 1.000 |
| **Waist Circumference, mean (SD)** |  | 0 | 99.4 (19.3) | 114.1 (15.1) | 84.2 (7.9) | <0.001 |
| **Hip Circumference, mean (SD)** |  | 0 | 108.8 (17.5) | 121.8 (14.2) | 95.4 (7.4) | <0.001 |
| **BMI, mean (SD)** |  | 0 | 30.1 (8.4) | 37.3 (5.5) | 22.6 (1.7) | <0.001 |
| **Systolic Blood Pressure, mean (SD)** |  | 12 | 127.8 (22.0) | 134.0 (19.5) | 121.2 (22.8) | 0.003 |
| **Diastolic Blood Pressure, mean (SD)** |  | 11 | 83.7 (14.9) | 87.8 (15.8) | 79.3 (12.7) | 0.004 |
| **Fasting Blood Sugar, mean (SD)** |  | 122 | 52.7 (60.4) | 49.2 (59.9) | 58.2 (62.4) | 1.000 |
| **Respiratory Rate (cpm), mean (SD)** |  | 24 | 19.8 (2.3) | 20.0 (2.4) | 19.6 (2.2) | 1.000 |
| **Pulse Rate (bpm), mean (SD)** |  | 19 | 82.2 (12.7) | 82.8 (12.7) | 81.6 (12.8) | 1.000 |
| **Temperature (Celsius), mean (SD)** |  | 12 | 36.5 (0.3) | 36.5 (0.3) | 36.5 (0.3) | 1.000 |

**Figure 1.**
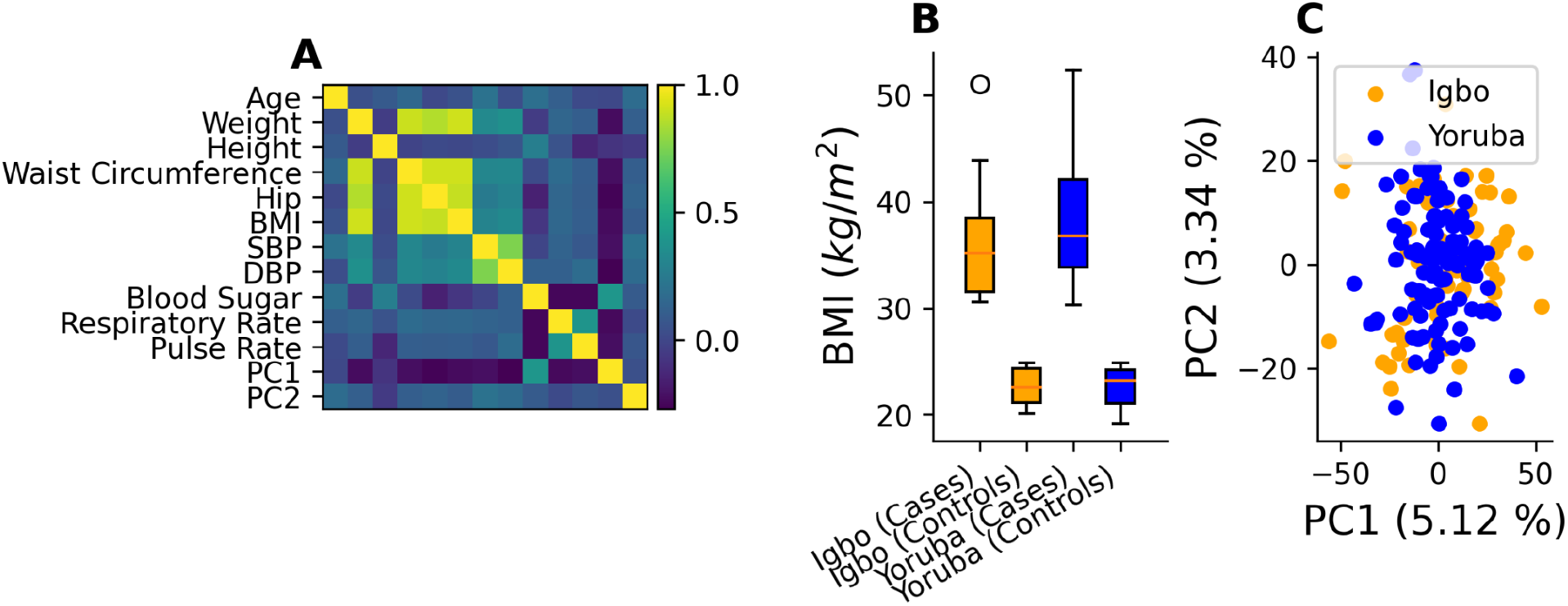
**(A)** Correlation across quantitative phenotypes for 171 post-QC subjects for this study. Note that anthropometric traits appear tightly correlated, as expected. **(B)** Distribution of BMI across Yoruba and Igbo ethnolinguistic groups stratified by case and control status **(C)** Top principal components computed using proteomic variation do not notably distinguish between ethnolinguistic groups.

We assessed how phenotypic and proteomic variation vary across the two ethnolinguistic groups sampled in our study, the Igbo and Yoruba groups from Nigeria. During the study design process, we sought to balance the distribution of cases and controls across both ethnolinguistic groups (see Methods). This ensures that we observe that there are no statistical differences in BMI across the ethnolinguistic groups (p = 0.49; Kolmogorov-Smirnov Test, Figure 1B). We also investigated whether proteomic variation explains substantial differences between the self-identified ethnolinguistic groups (Figure 1C). Qualitatively, there appears to be little variation in protein abundances associated with ethnolinguistic groups.

### Differential Protein Expression between Obesity Cases & Controls

We used a logistic regression model to identify the proteins that are most differentially expressed between individuals with BMI > 30 and BMI < 30 (i.e. obesity cases versus controls) (see Methods). The model identified several proteins that exhibited significant differences in expression between cases and controls (Figure 2A).

**Figure 2.**
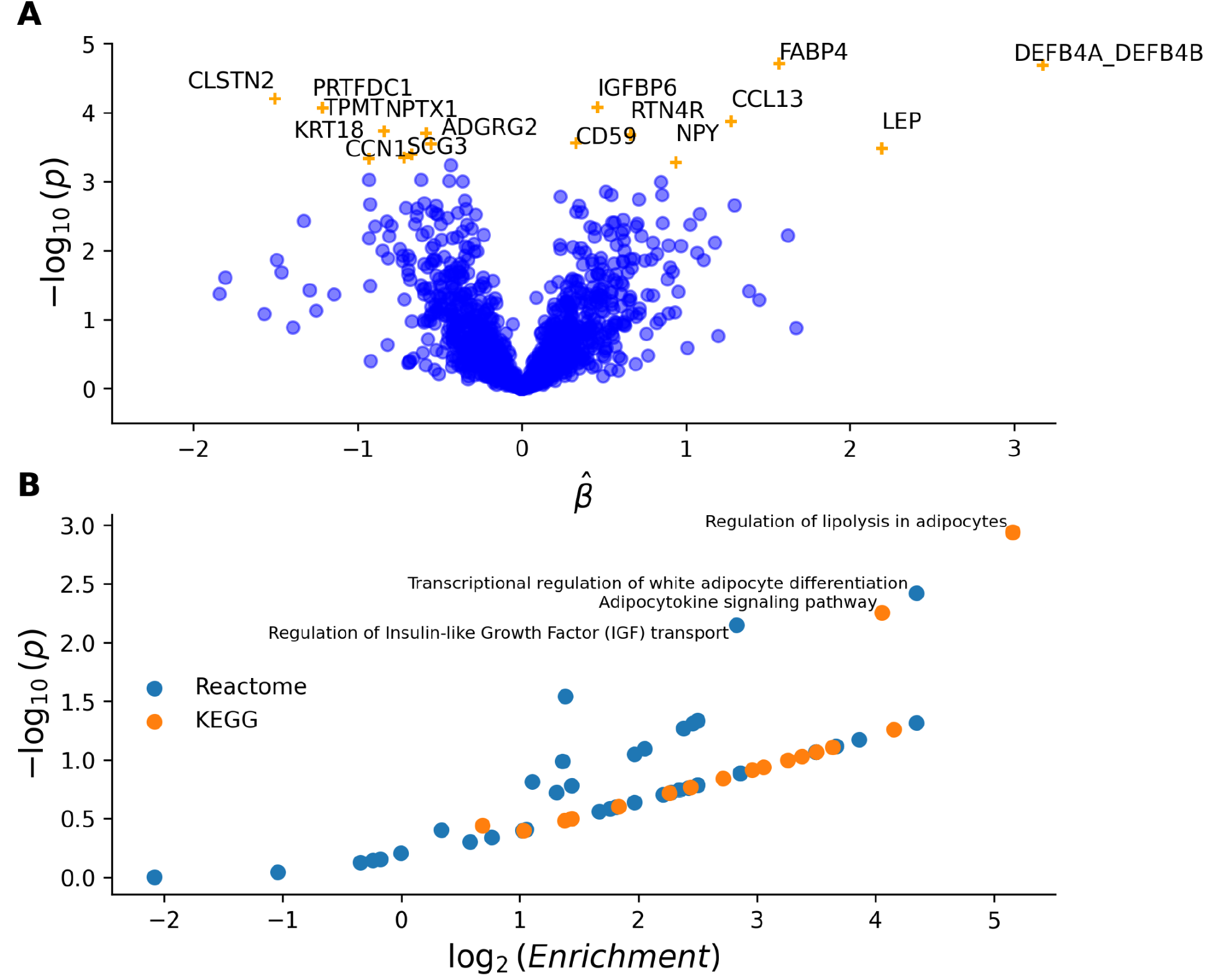
**(A)** Linear model of differential expression between cases and controls for obesity. Assays with significant differences between cases and controls at FDR < 0.05 are indicated in orange. Positive effects indicate a higher abundance in cases (e.g. BMI > 25) over controls. **(B)** Enrichments of pathways impacted by protein targets differentially expressed based on protein targets differentially expressed at an FDR < 0.05.

A total of 16 protein detection assays showed a significant difference between cases and controls (Figure 2A, Table S3). We find the results tend to cluster in three broad functional domains: adipocyte-related, inflammation, and neuropeptide-related effects on obesity. Adipocyte-related biological functions include a number of expected results, for example, proteins such as LEP and FABP4 which are well-established biomarkers of obesity and are elevated among high BMI individuals [17–22]. Indeed these are some of the strongest signals found within our dataset and also drive much of the observed pathway enrichments (Figure 2B).

Inflammatory responses to obesity have been well-established across both biomarker-based and clinical studies [23–26]. In our study, several inflammation and immune-response related protein biomarkers were found to have higher abundance in obesity cases relative to controls, such as DEF4A, DEF4B, CCL13, and CD59 (Figure 2A). One of the strongest signals, CCL13, has previous evidence of serum levels of CCL13 (C-C motif chemokine 13) associated with anthropometric traits and metabolic phenotypes, such as insulin and HDL-cholesterol in a Japanese cohort [27]. *CCL13* also has genetic variants associated with the pathophysiology of obesity in Hispanic children [28], lending additional evidence of its biological role in obesity-related traits.

We also found *SCG3* (Secretogranin 3) to be reduced in controls relative to cases, while *NPY* (Neuropeptide Y) was found to be more abundant in cases (Figure 2A, Table S3). The SCG2/SCG3 system is involved in the formation of secretory granules with appetite-related neuropeptides such as *NPY*, orexin, and *POMC* (Pro-opiomelanocortin) [29,30]. *POMC* is implicated in a monogenic form of early onset obesity and in populations of European ancestry, variants in *POMC* are associated with high BMI [31–34]. Two variants, rs16964465 and rs16964476, have been reported to affect the transcriptional activity of SCG3 and the minor allele containing haplotype is protective for high-obesity (odds ratio, 9.23; p=7 × 10^−6^) [30]. Notably, both of these variants are absent in European-ancestry groups but are common and in strong linkage disequilibrium in African and East-Asian ancestry groups (r^2^ = 0.993 and r^2^=1.00 respectively) [35]. The role of *NPY* in obesity has been described extensively in animal models, *in vitro* studies, and observational association studies [36–39]. Genetic associations between BMI and *NPY* in individuals of European ancestry (rs4307239, p=1 × 10^−11^), and meta-analyses have found variants in NPY to have bi-directional effects on obesity risk, suggestive of a larger tolerated range of NPY [13,40]. Indeed, small molecules such as *Velneperit* which inhibits *NPYR5*, an *NPY* receptor, have been established as proof-of-concept appetite regulators [41].

We also explored pathway enrichment from proteomic signals, to assess whether protein-derived risk factors are concentrated in specific biological pathways (see Methods). We observed that pathways related to adipocyte lipolysis, adipocytokine signaling, and insulin-like growth factors all were enriched among the significantly differentially expressed proteins (Figure 2B). These pathway enrichments are largely driven by the differential expression signals at FABP4, LEP, and NPY. Due to the smaller size (N=16) of the significantly differentially expressed gene-set, none of the enrichment results are statistically significant following correction for multiple testing (Table S4). Nonetheless, we take these results as suggestive that at a larger sample size, these or similar pathways would likely be enriched.

We also identified a group of proteins with no known genetic associations to BMI, obesity, or lean body mass in our study: *DEF4A_DEF4B, RTN4R, IGFBP6, CD59, ADGRG2, CCN1, TPMT, KRT18*, and *CLSTN2*. Some of these targets, such as CCN1 and IGFBP6, have some evidence of association with obesity-related phenotypes in cellular and animal models [42–46].

However, these broader signals merit further attention from larger-scale proteomic studies to validate their role in obesity pathogenesis.

### Protein QTL Detection

We conducted a scan for cis-pQTL (Protein Quantitative Trait Loci) to better understand the link between genetic variants and measured protein levels. Overall, we identified 73/1458 protein targets with significant sentinal cis-pQTL signal at an FDR < 0.05 (Table S5). These signals were isolated after performing conditional stepwise regression to assess whether multiple variants in *cis* to a protein target could explain the signal [16]. We also accounted for a number of covariates to ensure that these are free from effects due to obesity status, age, and other relevant covariates (see Methods).

Next we assessed the extent that allele frequency differences affect power for cis-pQTL discovery across ancestries. The ancestries sampled in this study are under-represented with regards to studies of functional genomic variation (specifically proteomic data), and we hypothesize that differences in variant frequency between our study and previously well-studied ancestries can substantially impact discovery power. For each of the 73 sentinel cis-pQTL variants, we investigated their frequency distributions across multiple worldwide populations, using GnomAD and 1000 genomes reference populations [47,48] (Table S5). Many of the variants we identified have a higher allele frequency in African-ancestry populations relative to non-Finnish European ancestry populations (Figure 3A). Of the 73 independent cis-pQTL variants detected, 21 were completely absent in European-ancestry populations, highlighting how leveraging diverse populations can lead to improvements in our understanding of genetic determinants of protein abundance. To quantitatively evaluate the impact of allele frequency differences across variants that are shared between ancestry groups, we looked at the relative sample size - the percentage increase in sample-size required to match statistical power between studies (see Methods). When looking at the relative sample size, we found that 31 of the 73 variants required sample size of greater than 10x in order to match the same power in a non-Finnish European ancestry population to our study based on allele frequency differences (e.g. *CLEC7A* at 355x and *KIR3DL1* at 250x) (Figure 3B, Table S5).

**Figure 3.**
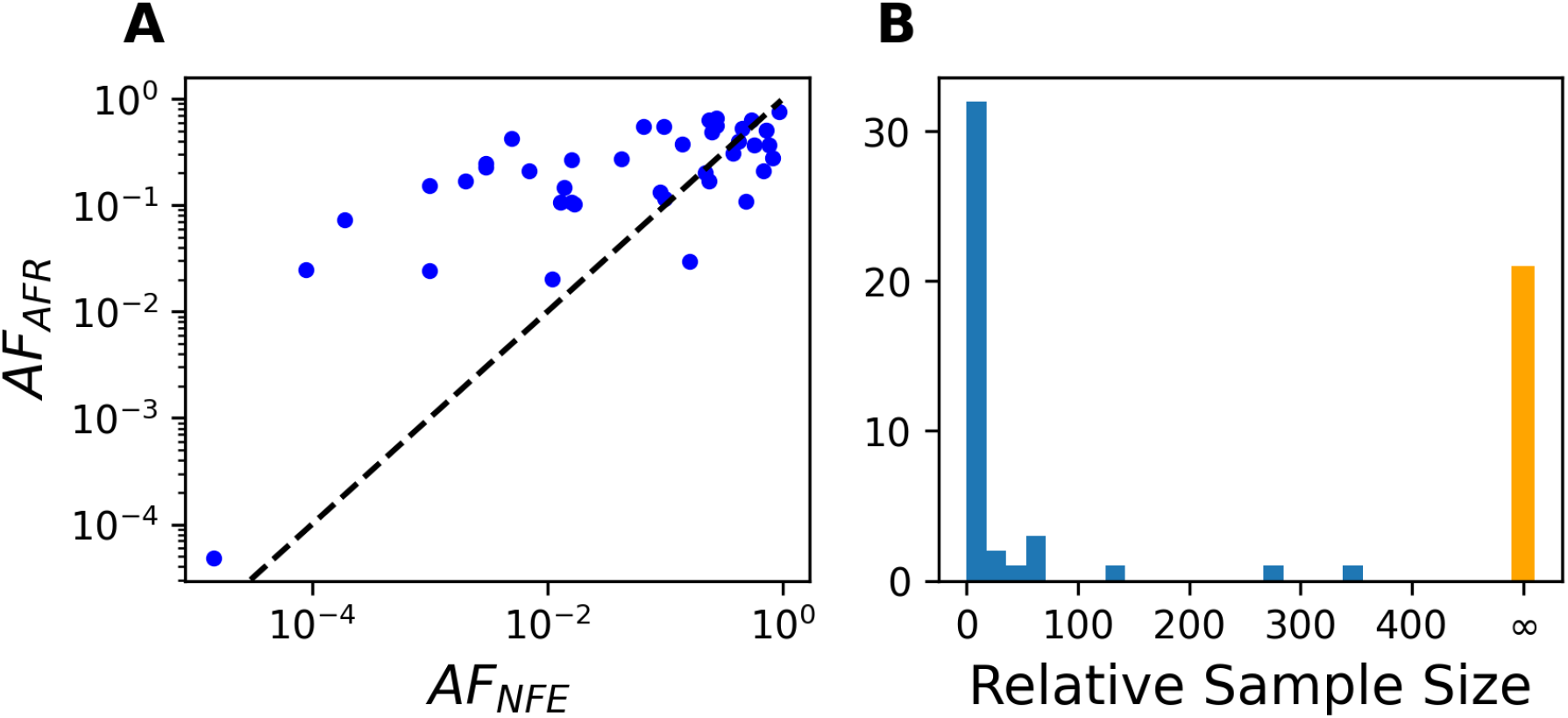
**(A)** Allele frequency differences at significant sentinel cis-pQTL variants between African population groupings in 1000 Genomes and GnomAD and Non-Finnish European populations, where the allele is observed in both ancestry groupings **(B)** Relative sample size required to detect the same cis-pQTL signal in Non-Finnish European ancestry populations, a measure of relative power for the specific sentinel allele detected. Note that the orange bar denotes sentinel variants that are detected *only* in African-ancestry populations, and hence have infinite relative sample-size.

For all of the significantly detected genes with a cis-pQTL in our experiment, we detected a single causal variant at an FDR < 0.05 for a specified protein target. This means that our experiment was well powered enough to detect a single causal variant, but no other variants within the cis-window are significantly associated with protein expression after conditioning on the lead variant. While allelic heterogeneity has been shown to be common across molecular traits, we suspect that at the current sample sizes we are likely only well-powered to detect the strongest effects [4,49]. We did not search beyond the cis-region of a protein target for trans-pQTL, reasoning that we were underpowered to detect notable trans-pQTL effects [4].

However, we do find several sentinel effects (N=21/73) at the cis-pQTL level where the lead variant is absent in non-Finnish European ancestry individuals in GnomAD (Figure 3B). To corroborate these findings we also leveraged a large dataset of cis-pQTLs from African-American ancestry individuals from the ARiC Study [6]. Of our 73 sentinel pQTLs, we found that 33 have a marginal p-value < 1e-3 in the ARiC Study, suggesting a modest rate of replication within African ancestry individuals (Table S5). Notably, of the 21 variants not found in non-Finnish European ancestries from GnomAD, only 4 of these variants have marginal p-values < 1e-3 in the ARiC dataset. Strikingly, 12 of these 21 variants do not have a marginal association at all in the ARiC proteomic dataset, likely due to their low frequencies in African -ancestry populations (mean allele frequency 0.03). While this suggests a modest rate of replication of these primary signals that we have observed, there are still several signals that are found in this dataset that are not well-explained in a larger dataset of African-American ancestry individuals (see Discussion).

There are a number of cis-pQTL that are strongly associated in our dataset, specifically for variants that are higher frequency in African populations (where we have higher statistical power). We found significantly associated variants regulating SIGLEC9 (rs273688) and CD33 (rs2455069), which are both immunoglobulin-like lectin family proteins (Figure 4A). While the variant rs2455069 for CD33 protein expression does not have a significant marginal p-value in the ARiC data, it is quite strongly associated in our data (see Discussion and Table S5). CD33-related sialic-acid-binding immunoglobulin-like lectin (SIGLECs) are inhibitory signaling molecules expressed on most immune cells and are thought to down-regulate cellular activation pathways via cytosolic immunoreceptor tyrosine-based inhibitory motifs [50]. The derived allele more common in African populations (rs273688) is a missense variant, which decreases the protein expression of SIGLEC9 in plasma; we hypothesize this may be in response to pathogen burden and the evolutionary compensation of the antiviral response [51]. While these two proteins are in the same gene family, it is notable that CD33 has an approved monoclonal antibody *Mylotarg* for acute myeloid leukemia (AML) [52]. In a study of clinical outcomes for European-ancestry AML patients, the presence of rs2455069 did not significantly alter clinical outcomes [53]. SIGLEC9 has also been proposed to promote immune-mediated escape from NK-cells, and knowledge of rs273688 reducing the abundance of SIGLEC9 could be valuable when determining the efficacy or dosage of CD33-targeting immunotherapies [51,54]. However, to our knowledge SIGLEC9 has not been in any drug target programs and warrants further inquiry into the evolutionary history of this high-frequency missense variant which reduces the abundance of this relevant immunological protein.

**Figure 4.**
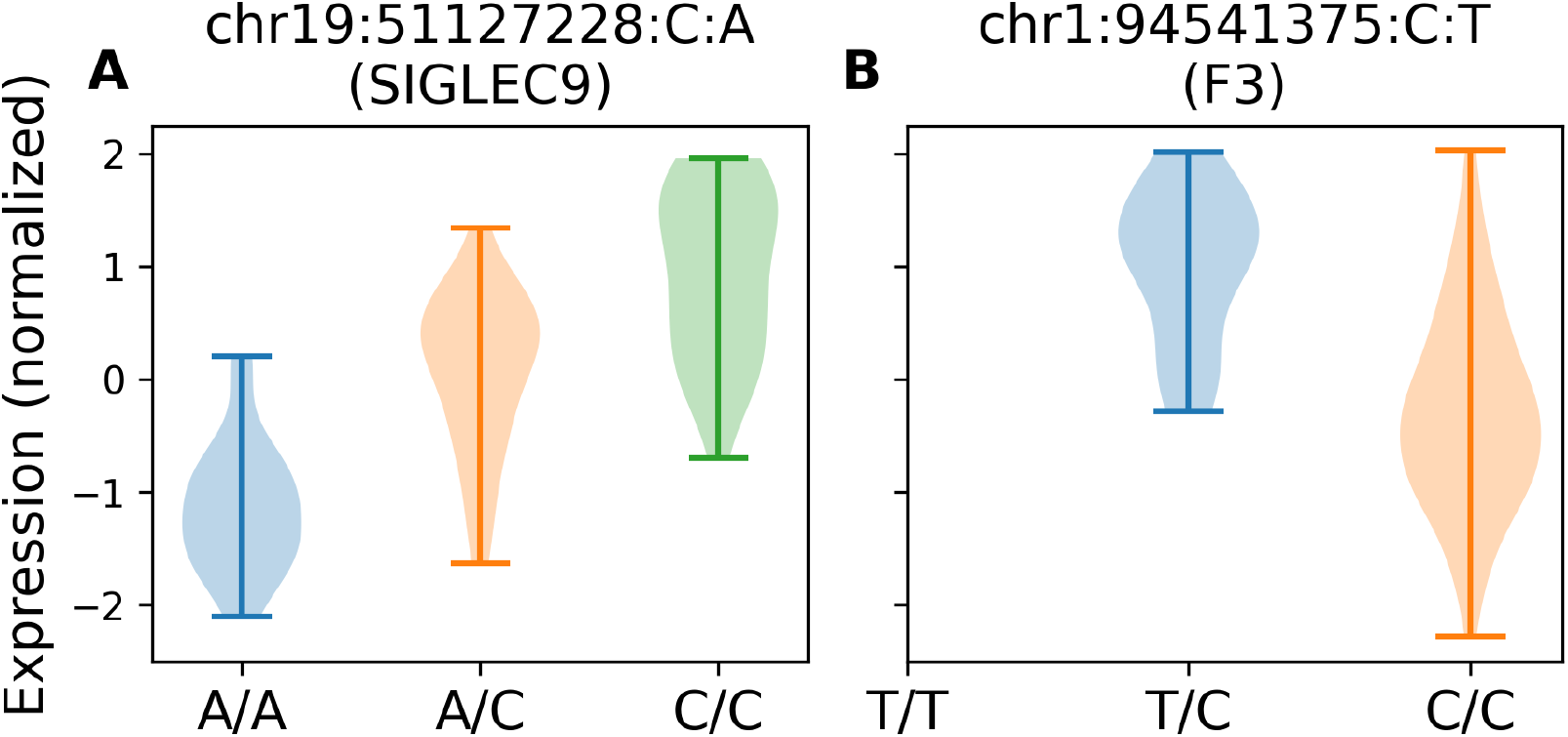
**(A)** Expression profile of sentinel missense variant and its influence on protein expression of *SIGLEC9*. **(B)** Sentinel variant that increases protein expression of *F3* (given sample sizes and the allele frequency of the variant, we did not observe any T/T homozygotes in our data)

We also found two variants associated with protein abundance of two coagulation factors, F3 and F7. Both of these variants (rs116174653 and rs564926165) are not found in non-Finnish European ancestry individuals from GnomAD and are not significantly associated in the larger ARiC dataset (Figure 4B, Table S5). Interestingly the effects of the variants (beta = 1.29 and beta = -1.92 respectively for rs116174653 [F3] and rs564926165 [F7]) are in opposite directions with respect to the derived alleles. From a pathway perspective, F7 is upstream of F3 in the cascade of platelet activation and thrombin production [55]. The variant rs564926165 associated with F7 protein abundance is actually a frameshift variant in *ADPRHL1*, which reverses ADP-ribosylation, which could potentially impact protein quantification of F7 (e.g. by reduction of antibody specificity) [56] (Table S5; see Discussion). Further research is required to determine the mechanism by which this variant may affect protein abundance measured through specific assays as a potential source of experimental noise.

Coagulation factors have a strong influence on the efficacy of blood-thinning medications, which are common treatments for thrombosis, hemorrhagic disease, and stroke [57]. Pharmacogenomic variants related to warfarin efficacy in individuals of African ancestry have been well-studied, specifically variants in *VKORC1, CYP2C9, CYP4F2;* although less variation in drug response is explained by these genic variants in individuals of African ancestry *[58,59]*. F7 has been associated with prothrombin time, and variants in F3 are associated strongly to fibrin-D dimer levels, a common measure of activated coagulation in individuals of European and African-American ancestries [7,8,60]. Fibrin-D dimer levels are highly predictive of coronary heart disease and stroke incidence, particularly in individuals of African ancestry [61]. These studies suggest that the marked increase in F3 protein abundance for carriers of the rs116174653 derived allele may warrant further study for its implications on pharmacological interventions using blood thinners.

**Table 2.**
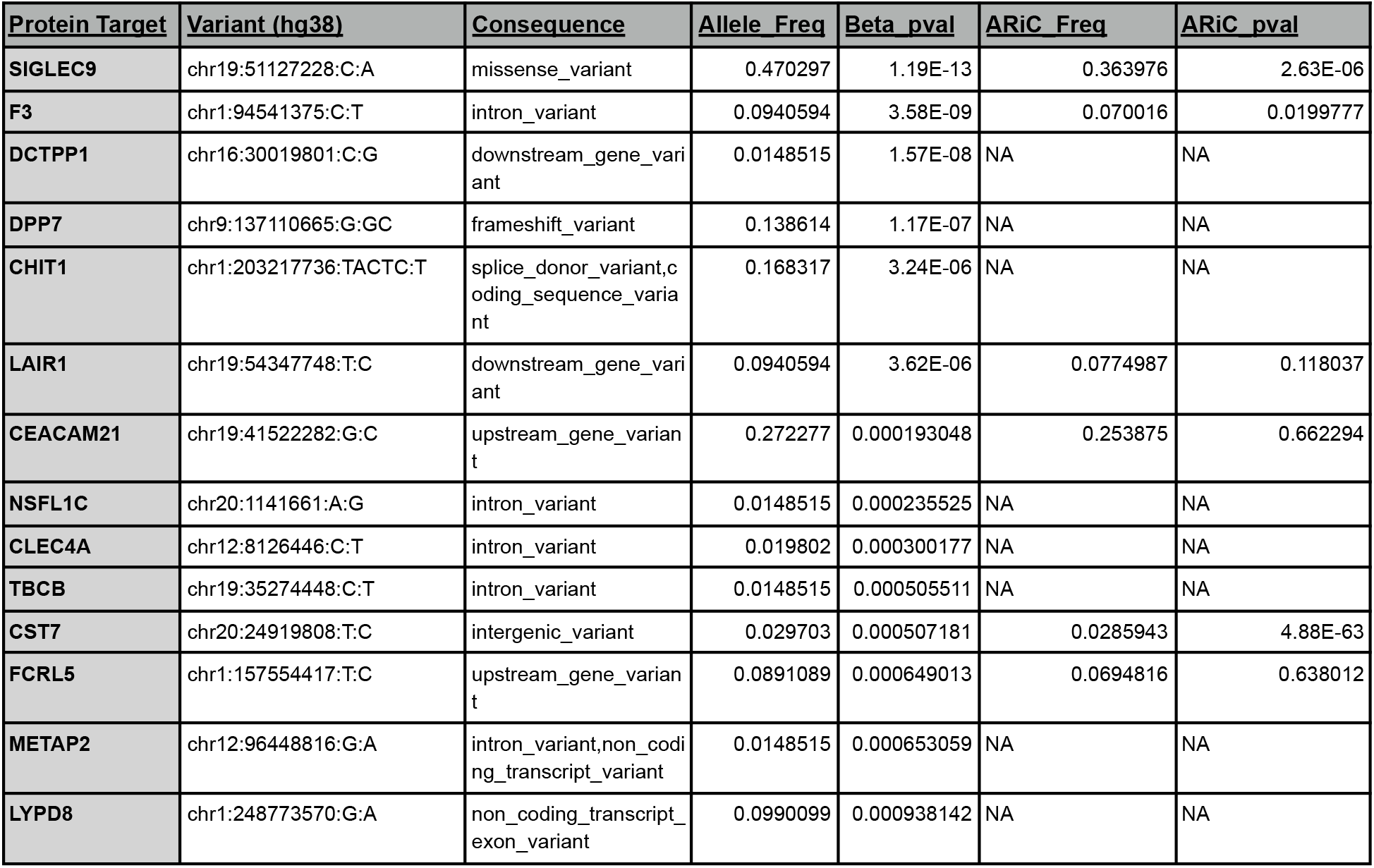
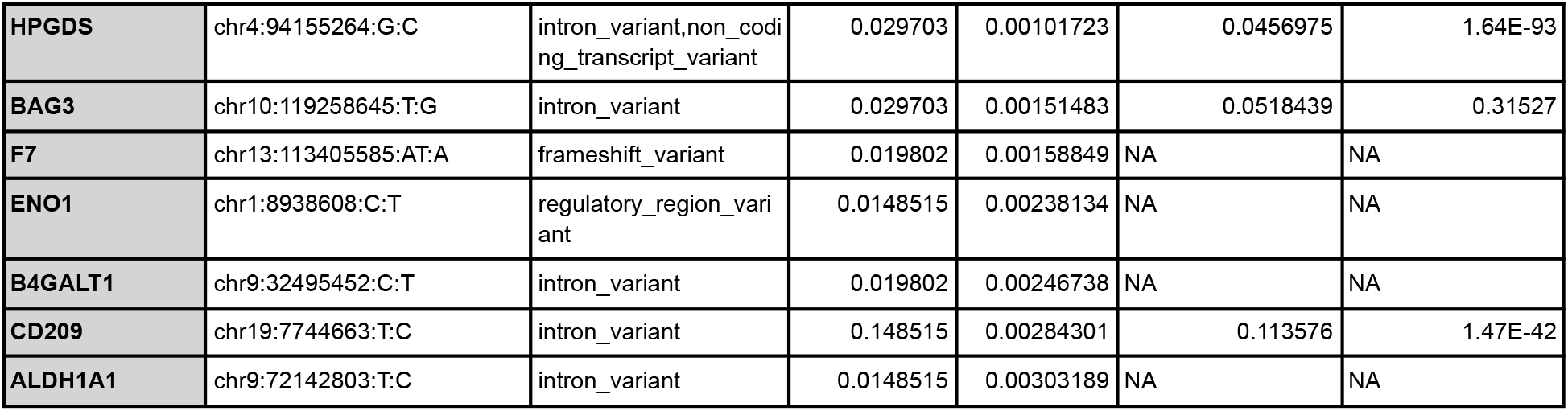
Significant cis-pQTL associations where variant is not found in GnomAD non-Finnish Ancestry individuals (N=21). Variant frequency in the ARiC dataset is also reported for variants whose summary statistics are available [6].

| Protein Target | Variant (hg38) | Consequence | Allele_Freq | Beta_pval | ARiC_Freq | ARiC_pval |
| --- | --- | --- | --- | --- | --- | --- |
| SIGLEC9 | chr19:51127228:C:A | missense_variant | 0.470297 | 1.19E-13 | 0.363976 | 2.63E-06 |
| F3 | chr1:94541375:C:T | intron_variant | 0.0940594 | 3.58E-09 | 0.070016 | 0.0199777 |
| DCTPP1 | chr16:30019801:C:G | downstream_gene_variant | 0.0148515 | 1.57E-08 | NA | NA |
| DPP7 | chr9:137110665:G:GC | frameshift_variant | 0.138614 | 1.17E-07 | NA | NA |
| CHIT1 | chr1:203217736:TACTC:T | splice_donor_variant, coding_sequence_variant | 0.168317 | 3.24E-06 | NA | NA |
| LAIR1 | chr19:54347748:T:C | downstream_gene_variant | 0.0940594 | 3.62E-06 | 0.0774987 | 0.118037 |
| CEACAM21 | chr19:41522282:G:C | upstream_gene_variant | 0.272277 | 0.000193048 | 0.253875 | 0.662294 |
| NSFL1C | chr20:1141661:A:G | intron_variant | 0.0148515 | 0.000235525 | NA | NA |
| CLEC4A | chr12:8126446:C:T | intron_variant | 0.019802 | 0.000300177 | NA | NA |
| TBCB | chr19:35274448:C:T | intron_variant | 0.0148515 | 0.000505511 | NA | NA |
| CST7 | chr20:24919808:T:C | intergenic_variant | 0.029703 | 0.000507181 | 0.0285943 | 4.88E-63 |
| FCRL5 | chr1:157554417:T:C | upstream_gene_variant | 0.0891089 | 0.000649013 | 0.0694816 | 0.638012 |
| METAP2 | chr12:96448816:G:A | intron_variant, non_coding_transcript_variant | 0.0148515 | 0.000653059 | NA | NA |
| LYPD8 | chr1:248773570:G:A | non_coding_transcript_exon_variant | 0.0990099 | 0.000938142 | NA | NA |
| **HPGDS** | chr4:94155264:G:C | intron_variant,non_coding_transcript_variant | 0.029703 | 0.00101723 | 0.0456975 | 1.64E-93 |
| **BAG3** | chr10:119258645:T:G | intron_variant | 0.029703 | 0.00151483 | 0.0518439 | 0.31527 |
| **F7** | chr13:113405585:AT:A | frameshift_variant | 0.019802 | 0.00158849 | NA | NA |
| **ENO1** | chr1:8938608:C:T | regulatory_region_variant | 0.0148515 | 0.00238134 | NA | NA |
| **B4GALT1** | chr9:32495452:C:T | intron_variant | 0.019802 | 0.00246738 | NA | NA |
| **CD209** | chr19:7744663:T:C | intron_variant | 0.148515 | 0.00284301 | 0.113576 | 1.47E-42 |
| **ALDH1A1** | chr9:72142803:T:C | intron_variant | 0.0148515 | 0.00303189 | NA | NA |

## DISCUSSION

The primary aim of our study was to investigate proteins whose expression were strongly associated with obesity status across the two ethnolinguistic groups in Nigeria. We were able to recapitulate well-known protein biomarkers of obesity (*LEP* and *FABP4*) and overall enrichment of adipocyte-specific biology. More broadly, we also found evidence of inflammatory markers (*CCL13*) and neuropeptides (*NPY*) differentiating obesity cases and controls, highlighting the multitude of biological systems contributing to and affected by obesity. While we were only well-powered to detect only the strongest effects due to limited sample size (N=176), we expect that a larger study with similar targeted proteomics could support further discovery.

A secondary aim of our study was to highlight genetic variants in African-ancestry populations that regulate proteomic variation. Increasingly, genetic regulation of protein targets have been utilized to understand the degree to which genetic effects are mediated by protein abundances [4,6,16]. However, to the best of our knowledge there have been few proteomic studies in individuals of African ancestry (the study of [6] is a notable exception that has a large sample of African-American ancestry individuals). African-sample proteomic studies are critical to enable comprehensive prediction of protein targets from genetic data, specifically for variants that are rare outside of Africa. In our work, even with modest sample sizes (N=101), we found a sizeable fraction (21/73) of significant cis-pQTL that were absent in non-African catalogues of genetic variation.

While many of our pQTL variants were replicated in the larger analysis of the ARiC study (33/73), there are still a substantial number that were not reliably replicated. There are several potential reasons for this lack of replication, each with notable solutions. The first is simply that the tested allele was absent in the African-American ancestry individuals of the ARiC proteomics study, motivating the development of proteomic cohorts from continental Africa particularly for replication of rarer variant effects [6]. A second reason is that we utilized the Olink Explore 1536 platform and the ARiC study analyzed data from the Somalogic v4 platform, suggesting differences in proteomic assays can mediate replication [6]. The correlations in protein measurements between these two technologies have been investigated in great detail, with the current established consensus that each of these assays vary substantially in their relative protein quantification measurements [62,63]. Therefore, we expect that applying a wider array of proteomic assay types may contribute to additional pQTL replication.

As genomic and proteomic sampling efforts increase in Africa, we expect additional pQTLs will be found (both *cis* and *trans* acting variants), improving their reliability for downstream analyses such as Mendelian Randomization and proteome-wide association studies (PWAS) [5]. More importantly, as specifically highlighted for the novel cis-pQTL of *F3*, it is an open question as to the value of leveraging targeted proteomic information when making predictions on pharmacogenomic intervention. We expect that these assays are more reflective of a more direct endpoint of the molecular cascade, potentially leading to more actionable evidence for drug dosage and administration.

In this work, we highlight the relative importance and relevance of the two distinct aims of this project: (1) elucidating proteomic differences between obesity classes, an NCD of increasing prevalence on the African continent and (2) genetic variants that influence protein products within African populations. The former contributes to a deeper molecular perspective on an increasingly prevalent health condition in Africa, highlighting potential avenues for preventative intervention and monitoring. The latter contributes to our understanding of genetic variants which affect protein abundance in plasma, opening up possibilities for robust genomic analyses and improving the portability of predictive models to African-ancestry individuals.

## METHODS

### Sample Enrollment, Consent and Biobanking, and Phenotype Collection

We recruited a total of N=176 individuals according to the following inclusion criteria: (1) participants who voluntarily provide Informed consent for the study and (2) participants aged 18 to 90 years. We excluded any individual who withdrew consent at any point in the study and individuals without the capacity to provide consent (or have a legal guardian provide consent).

Samples were collected from three different sites: Abeokuta Federal Medical Centre (ABK), University of Nigeria Teaching Hospital (UNTH), Ebute Metta Federal Medical Centre (EBM). These are all tertiary medical centers across three different states in Nigeria (Ogun, Enugu, and Lagos). Ethical approvals were obtained from the IRB at each of these sites.

Phenotypes were collected electronically using SurveyCTO and subsequently harmonized using a custom in-house process-phenotypes R package (see Resources). Questions were organized broadly to collect demographic information, medication history, and phenotypic information on obesity and associated co-morbidities. Using the initial phenotypes collected, propensity score matching was performed to ensure that cases and controls defined by BMI > 30 kg/m^2^ and BMI < 30 kg/m^2^ respectively, are matched according to age, sex, and ethnolinguistic group labeling. This ensures that each case has a similarly sampled control with respect to external metadata and that there were an equal number of cases and controls (Figure 1B).

### Lab processing

Whole blood aliquots from samples were biobanked at -80°C ahead of DNA extraction and genotyping while the plasma aliquots were re-spun for further processing ahead of protein testing. DNA from whole blood samples were extracted using the MagMax Blood/Saliva Extraction kit on the Kingfisher Nucleic Acid Extraction System. Quantification of DNA was measured using fluorescence intensity of Quantifluor One dsDNA dye (Promega Quantifluor One dsDNA dye; E4871), inside a 0.5 milliliter thin-walled PCR tube. The Quantifluor One dsDNA dye specifically binds to double-stranded DNA and the dye fluorescence can be measured as a quantitative representation of the concentration of DNA present in the dye solution. Samples within the tolerable range of 5-10 nanograms of total DNA (typical yield for 0.3-0.4 milliliters of whole blood as starting material) were submitted for genotyping.

Genotyping was performed using the GeniiSys 1.0 Array on an Illumina iScan machine [[64]. Plasma samples for proteomic evaluation were sent to Olink Proteomics in Waltham, MA and processed according to best practices for the Olink Explore 1536 assay including establishment of internal controls for quantification of protein abundance using an antibody-based protein extension (PEA) assay.

#### Quality Control

##### Phenotype Quality Control

Phenotypes were collected using SurveyCTO (see Sample Enrollment Section), and subsequently processed using the `54gene-phenotype-processing` workflow (see Resources) using form definitions downloaded on July 28, 2022. All phenotypes were standardized to have the same unit (Table 1). Two individuals (from N=176) were excluded due to > 50% phenotype-missingness across the set of phenotypes that were collected (Table 1) leaving us with N=174 individuals passing phenotype-level quality control.

##### Proteomic Quality Control

To reliably control for extraneous sources of variation in Olink Explore 1536 proteomics data, we developed a well-tested and automated quality control pipeline to reduce the influence of outliers and extraneous sources of variation (see Resources). Since the units of analysis are Sample x Panels pairs for Olink proteomics data, each sample is run on each panel pertaining to proteins for a specific disease area, and we filter these combinations for outliers specifically.

In addition to exclusions applied from the previous phenotyping quality control section that removed two individuals, we applied the following quality control thresholds:

1. Filter out assays that have an “Assay Warning” indicator from Olink
2. Filter out Sample x Panel combinations that are:

a. > 3 standard deviations away from the median NPX value
b. > 3 standard deviations from the mean PC1 or mean PC2 based on NPX values per-sample
c. > 3 standard deviations from the interquartile range based on NPX values

In total, our quality control procedure removed 6 assays due to Assay Warnings (Supplementary Table 1) and 24 Sample x Panel exclusions (Supplementary Table 2). This removed a further 3 individuals entirely (all panels from these individuals were removed as outliers) from the analysis leading to 171 individuals that are carried forward for analyses of proteomic differential expression.

##### Genotype Quality Control

All samples were genotyped using our custom Illumina GeniiSys v1.0 Array [64] and imputed using the TOPMed imputation server, while filtering to variants in minor allele frequency bins to maintain a mean imputation r^2^ > 0.9 in each bin [65]. For initial genotyping quality control, the following steps were applied: (1) filtering to MAF > 1% and genotype missingness < 5%, (2) subset to subjects per-chip that are at less than 3^rd^ degree relatives [66], (3) remove samples that have discordant self-reported and genetic sex. Overall, we retained 101 samples for all analyses with linked proteomic data for downstream cis-pQTL evaluation.

#### Linear Modeling

When evaluating the contribution of proteomic variation to differences between obesity cases and controls we employed a logistic regression framework to evaluate the association between the normalized protein abundance and the log-odds of being a case (BMI > 25). We accounted for self-identified ancestry group, sampling site, sex, age, systolic blood pressure, fasting blood sugar, and 5 proteomic-based PCs as covariates within these models. To correct for multiple testing we established an FDR < 0.05 using the q-value method from [67].

#### Pathway Enrichment Analysis

We used Webgestalt (WEB-based Gene SeT AnaLysis Toolkit) [68] to carry out functional enrichment analysis of differentially expressed genes identified in our analysis, intersecting KEGG and Reactome gene sets [69,70]. Since the proteins included on the Olink Explore 1536 assay contains a targeted set of proteins across four different disease domains, we accounted for the full set of proteins as the background distribution when conducting enrichments to avoid the effect of protein ascertainment. When evaluating statistical significance of pathway enrichments for differentially expressed proteins we used an FDR < 0.05 (Table S7).

#### Protein Quantitative Loci Calling

For calling cis-pQTL we used the software TensorQTL [71]. We included the top five genotype-derived principal components, two proteomic-derived principal components, BMI, sex, age, and self-identified ancestral group as covariates into the model. Our primary goal was to avoid potential conflation of the effect with that of BMI, which was a major biological aim of the study. We employed a nominal false discovery rate (FDR) cutoff of FDR < 0.05 for each of the cis-pQTL signals and subsequently employed a conditional stepwise regression step, that allows for multiple signals within a locus [72]. In all cis-pQTL analyses we used the TensorQTL default of one megabase around the transcription start site of the gene target as the cis-window. Variant annotations were obtained using VEP release 108 [73]

#### Assessment of Relative Power for pQTL

When comparing the relative effective sample size we use the following expression for association mapping power based on simple linear regression [74]:

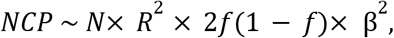

where NCP is the non-centrality parameter of a non-central chi-squared distribution with one degree of freedom, *N* is the sample-size, *R*^2^ is the imputation dosage quality measure, *f*is the allele frequency, and β is the true causal effect of the variant. When assessing relative-power, we hold all variables but *N* and *f* constant across two studies to equate their non-centrality parameters. This then leads to the following relationship (where study index is denoted by subscripts):

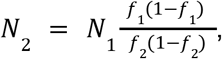

where *N*_2_ is the sample size-required for the second study to have equivalent power to the first study accounting for the difference in allele frequency. In Figure 3B, we present this simply as the ratio on the right-hand side of the above expression as that is the scalar factor by which the sample size must be increased to recover the same marginal effect (rather than the absolute number of samples required).

We have made the assumption that for imputed variants, the accuracy of imputation remains the same across allele frequencies, but this assumption is less tenable at lower frequencies [65,75].

This most acutely impacts the assumptions when *f*_2_≪*f*_1_(or the opposite case), but_*R*_^2^thresholding variants to have a high imputation *R*^2^will limit the impact of this on comparisons in power. Specifically, in the cases when *f*_2_≪*f*_1_, we expect *R*^2^_2_≪*R*^2^_1_, which should increase the sample-size required in the second study based on the equation above, suggesting that in this regime of joint allele frequencies we conservatively underestimate the sample-size required for equivalent power (Table S5).

## SUPPLEMENTARY TABLES

**Table S1:** Olink Explore 1536 assays removed due to “Assay Warning” tags indicating experimental measurement error.

**Table S2:** Sample x Assay combinations removed due to outlier removal procedures.

**Table S3:** Summary statistics of differential expression linear model. NOTE: effect direction here is opposite sign to that in Figure 2A due to coding of Case/Control status.

**Table S4:** Pathway enrichment statistics from KEGG & Reactome using WebGestalt.

**Table S5:** Independent cis-pQTL marginal summary statistics. Header columns with prefix

’gnomADg’ are allele frequencies across gnomAD genomes, ‘gnomeADe’ are allele frequencies across gnomAD exomes; Allele frequencies from 1000 Genomes regional groupings are also shown via an ‘AF’ suffix (e.g. ‘AFR_AF’). Relative sample-size is calculated using frequency of variants observed in EUR & AFR groupings from 1000 Genomes or GnomAD if the variant is not found in 1000 Genomes regional groupings. Additional column headers are derived from VEP.

## ACKNOWLEDGEMENTS

We would like to acknowledge all of the participants in this study. We also thank members of the facilities at 54gene Inc, Lagos (Nigeria), Olink, Waltham (USA) for genotyping and proteomic assay management respectively. We thank the site investigators and the 54gene site quality assurance officers for effectively coordinating the study at the various sites.

## DECLARATION OF INTERESTS

At the time research was conducted, A.B., K.P.d.A., G.E., C.N., Y.I., O.O., E.J, A.Y., A.E.O., J.P., C.O’D., and P.F. were employed by 54gene, Inc and are beneficiaries of compensation through this employment. Funding for this study was provided by 54gene, Inc.

## RESOURCES & SOFTWARE

1. Olink Quality Control Pipeline: https://gitlab.com/data-analysis5/proteomics/54gene-olink-qc
2. Phenotype Processing Package: https://gitlab.com/data-analysis5/phenotypes/process.phenotypes
3. TensorQTL: https://github.com/broadinstitute/tensorqtl
4. TopMED Imputation Server: https://imputation.biodatacatalyst.nhlbi.nih.gov/#

## DATA ACCESS AND AVAILABILITY

Imputed genotypes, phenotype, and proteomic data are available from the authors upon request. Summary statistics for differential expression analyses and associated cis-pQTL and are available in Table S3 and Table S4 respectively.

